# A general framework for species-abundance distributions: linking traits and dispersal to explain commonness and rarity

**DOI:** 10.1101/2022.04.15.488506

**Authors:** Thomas Koffel, Kaito Umemura, Elena Litchman, Christopher A. Klausmeier

**Affiliations:** W. K. Kellogg Biological Station, Michigan State University. <u>Email</u>; Graduate School of Human Development and Environment, Kobe University. <u>Email</u>; W. K. Kellogg Biological Station, Department Integrative Biology, Program in Ecology & Evolutionary Biology, Michigan State University. Department of Global Ecology, Carnegie Institution for Science. <u>Email</u>; W. K. Kellogg Biological Station, Departments of Plant Biology & Integrative Biology, Program in Ecology & Evolutionary Biology, Michigan State University. <u>Email</u>

**Author notes:** <u>Corresponding Author</u>: Thomas Koffel.

**Keywords:** trait-based model, species-abundance distributions, trait-abundance distributions, metacommunity, competition, mass effects

## Abstract

Species-abundance distributions (SADs) describe the spectrum of commonness and rarity in a community. Beyond the universal observation that most species are rare and only a few common, more-precise description of SAD shape is controversial. Furthermore, the mechanisms behind SADs and how they vary along environmental gradients remain unresolved. We lack a general non-neutral theory of SADs. Here we develop a trait-based framework, focusing on a local community coupled to the region by dispersal. The balance of immigration and exclusion determines abundances, which vary over orders-of-magnitude. Under stabilizing selection, the local trait-abundance distribution (TAD) reflects a transformation of the regional TAD. The left-tail of the SAD depends on scaling exponents of the exclusion function and the regional species pool. More-complex local dynamics can lead to multimodal TADs and SADs. Connecting SADs with trait-based ecological theory provides a way to generate more-testable hypotheses on the controls over commonness and rarity in communities.

## Introduction

Species-abundance distributions (SADs) describe the distribution of population densities of all the species in a community. They are intermediate-complexity descriptors of the diversity of a community: more informative than species richness but less detailed than a list of species and their abundances (McGill et al. 2007). Over almost a century, ecologists have collected countless SADs from natural communities. The universally observed pattern is that most species in a community are rare, while a few are common, with abundances ranging over orders of magnitude (McGill et al. 2007). Further generalities about the shape of SADs remain controversial (Ulrich et al. 2010), as do the mechanisms behind them. Common species drive ecosystem functioning (Grime 1998, Winfree et al. 2005), while rare species can provide adaptive capacity in changing environments (Yachi & Loreau 1999, Norberg et al. 2001) but face higher risk of extinction (Terborgh & Winter 1980). Thus, to understand the mechanisms behind SADs is to understand how communities are structured, which is essential to predict how they will reorganize under anthropogenic environmental change (e.g., climate change and eutrophication).

McGill & colleagues (2007) categorized 27 theoretical models of SADs, including purely statistical models (log-series, Fisher et al. 1943; log-normal, Preston 1948), nicheapportionment models (geometric, Motomura 1932, Doi & Mori 2013; broken stick, MacArthur 1960, Sugihara 1980, Tokeshi 1990), and single-species models (e.g., Engen & Lande 1996ab). However, only mechanistic models based on multispecies population dynamics can simultaneously generate predictions about how species’ abundances relate to their traits, how community structure changes along environmental gradients, and how communities will reorganize under environmental change, so we focus on them.

Vellend (2010) identified four key processes that shape communities: selection (niche differences), ecological drift (demographic stochasticity), speciation, and dispersal. Hubbell’s neutral theory (2001) combines ecological drift, speciation, and dispersal but lacks selection. It generates realistic SADs, but its central assumption that species are identical contradicts patterns found in nature (Harpole & Tilman 2006). However, with some notable exceptions (Engen & Lande 1996ab, Wilson et al. 2003), most purely niche-based models fail to generate realistic SADs: it can be difficult to get more than a few species to coexist (Edwards et al. 2018), and when they do, they typically have comparable abundances. It is unlikely that purely niche-based models can recreate the orders-of-magnitude variation observed in species abundances without carefully tuned parameters.

One possible solution to this conundrum is to incorporate processes acting at broader spatial or temporal scales into our explanations of local community structure. Spatial and temporal heterogeneity in selection provides a powerful mechanism of species coexistence (Chesson 2000, Mouquet & Loreau 2003). Immigration to a local community can occur in space, from other communities in the region (regional species pool, assumed to be fixed), or in time, from a reservoir of resting stages. The addition of immigration to niche-based models trivially solves the problem of the local co-occurrence of many species (Shmida & Ellner 1984, Loreau & Mouquet 1999), but as we will show, naturally results in order-of-magnitude variation in population density. Empirical studies have shown that such “mass effects” play an important role in maintaining local diversity (Shmida & Wilson 1985).

While simulations of some particular models have shown that combining immigration from a regional species pool with local interactions results in realistic SADs (Hughes 1986, Mouquet & Loreau 2003, Wilson & Lundberg 2004, Vergnon et al. 2012), they are not amenable to analytical treatment and hard to generalize. Therefore, a general theoretical framework that merges local, niche-based, and regional processes to explain SADs and can be applied to a wide range of questions, is lacking. In this paper we develop such a framework, which combines local interactions with immigration from a regional species pool to derive community structure. This theoretical framework exposes the essential elements of these previous models.

One barrier to general theories of ecological communities is the “curse of dimensionality”: as the number of species 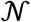 goes up, the number of parameters typically goes up as 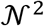. For realistically sized communities, this becomes unmanageable: there are more parameters than empiricists can measure and too many degrees-of-freedom for theoreticians to constrain their models. One theoretical solution is to choose parameters randomly *(e.g.,* May 1972, Wilson & Lundberg 2004, Barbier et al. 2018, Allesina & Grilli 2020), but this allows prediction only in an ensemble of replicate communities and is disconnected from the biology of how species interact with each other and their environment.

An alternative solution is to take a *trait-based* approach, where species are defined by the functional traits that determine their ecological performance (McGill et al. 2006, Litchman & Klausmeier 2008). The closely related theoretical frameworks of evolutionary game theory (Brown & Vincent 1987, McGill & Brown 2007) and adaptive dynamics (Geritz et al. 1998) provide elegant tools for formulating and analyzing trait-based models (reviewed in Klausmeier et al. 2020), but assume closed systems. While these trait-based approaches consider the interaction of a continuum of strategies and therefore have the potential to model diverse communities, the resultant communities are often species-poor unless extreme trade-offs are assumed (Edwards et al. 2018). Therefore we apply the results of our general dispersal-selection theory to trait-based models in a metacommunity setting, where local dynamics is coupled to a regional pool of species through dispersal. This trait-based approach generates a broader range of testable predictions beyond the shape of the SAD — the traits of common vs. rare species and how SADs vary along environmental gradients — which will enhance its falsifiability (McGill et al. 2007).

### General framework

Consider the general class of models of 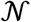 species, whose dynamics in a local community are given by

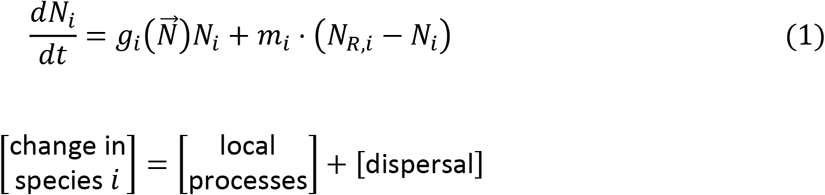

where *N_i_* is the density of species *i*. The first term describes local processes, with per capita growth rate *g_i_*, which depends on the vector of all species, 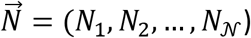. The second term models dispersal at rate *m_i_* between the local community and the broader region, where the species has fixed density *N_R,i_*. The equilibrium of eqn. (1) is typically analytically intractable, so it usually must be studied numerically. However, we can derive analytical results in the limits of small and large dispersal to guide our understanding.

We assume that the community goes to an equilibrium in the absence of dispersal (*m_i_* = 0) where each species either persists (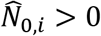 and *g_i_* = 0) or goes extinct (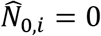 and *g_i_* < 0). We call species that persist in the absence of dispersal *(true) core species.* With even small dispersal, non-core species achieve positive abundance through mass effects. When they remain locally rare, we call them *satellite species* following Hanski (1982) (although we do not assume they are also regionally rare).

In the limit of small dispersal (*m_i_* ≈ 0), the abundance of core species is only marginally affected by dispersal. In contrast, the equilibrium abundance of satellite species is determined by the balance of the immigration rate *m_i_N_R,i_* and exclusion rate *e_i_* ≡ – *g_i_*. At the lowest order approximation,

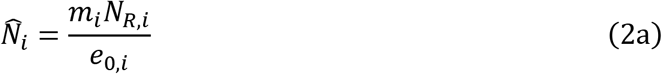

where *e*_0,*i*_ is the exclusion rate of species *i* in the absence of dispersal i.e., 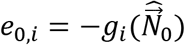.

Equivalently,

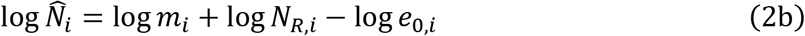

which suggests that equilibrium population sizes typically vary over orders of magnitude, as in empirical SADs. A slightly refined approximation accounts for emigration:

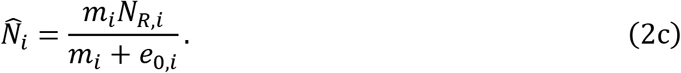

which shows that in the limit of large dispersal (*m_i_* → ∞), dispersal swamps local processes, so that each species simply matches its regional abundance: 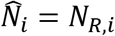.

The universal observation that SADs consist of few common species and many rare species leads us to hypothesize that, using the appropriate currency for abundance (Morlon et al. 2009), the common species represent core species and the rare species represent satellite species dependent on continued dispersal from elsewhere to persist. Given this hypothesis, the bulk of SADs consists of satellite species, whose abundances are determined by the balance of immigration and exclusion given by eqn. (2), while the core species’ abundances are determined mainly by local processes. Taking the dispersal rates *m_i_* and the regional abundances *N_R,i_* as given model parameters, the final ingredient that determines the SAD is the distribution of exclusion rates, *e_i_*. Therefore, we must specify how species interact, which is encoded in the growth functions 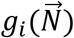 in eqn. (1).

If we had detailed information on the interactions within a community such that we could parameterize the growth functions 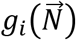 in eqn. (1), we could predict how its SAD depends on dispersal with the regional metacommunity. However, these predictions would be specific to that particular system with defined interactions, limiting generality; furthermore, such knowledge is typically lacking, particularly for species-rich communities. Therefore, to make general predictions about SADs, we need a reasonable way to parameterize models of diverse communities. Due to its solid basis in functional ecology (McGill et al. 2006, Violle et al. 2007, Litchman & Klausmeier 2008), we use *trait-based* ecological theory (Klausmeier et al. 2020) to reduce the dimensionality of parameter-space in the rest of this paper.

### Trait-based models

The fundamental assumption of trait-based models is that a species’ demographic rates in a given abiotic/biotic environment depend only on its traits, 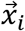. In this case, our general model (eqn. 1) becomes

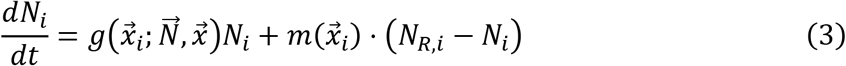

Note that the *growth function g* in eqn. (3) is not species-specific, but shared by all species in a guild (Brown & Vincent 1987). Therefore, species that have identical traits have the same per capita growth rate, rendering them ecologically neutral at the local scale. The growth function *g* encodes density-dependence and interspecific interactions, and depends on both the traits of the focal species, 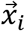, and the density and traits of the entire community, 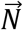 and 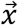. The traitbased framework of eqn. (3) is quite general. In the absence of dispersal, it has been used to model many ecological scenarios, including complexities such as spatial and temporal heterogeneity and population structure (see Klausmeier et al. 2020 for an overview). Thus, diverse ecology can be encoded in the deceptively simple-looking *g*. The dispersal rate *m* can also depend on species traits or be assumed constant. The model description is completed by the description of the regional species pool through their traits 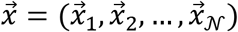 and abundances 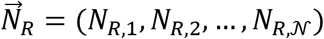.

To facilitate analysis, instead of considering a large set of discrete species, we take a continuum limit, where individuals are indexed by their trait values. Analogous to eqn. (3), the general form of such a continuum model is

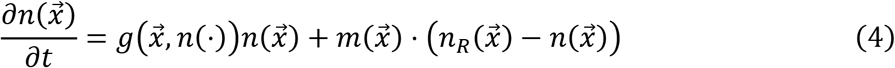

for the dynamics of the local population-density 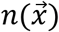 of individuals with trait values 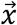, where *n*(·) denotes the *local trait-abundance distribution* (TAD). Note that 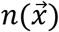 is a density per unit trait, and therefore differs from the *N* of particular species in eqn. (3). The regional species pool is described by the *regional TAD* 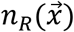 and the *species-density function* 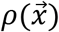, which describes how the species are distributed in trait-space (see Fig. 1C). The species-density function is essential for converting from this continuum approach back to discrete species, which constitute SADs (described in Box 1). Finally, 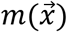 is the potentially trait-dependent dispersal rate.

**Figure 1.**
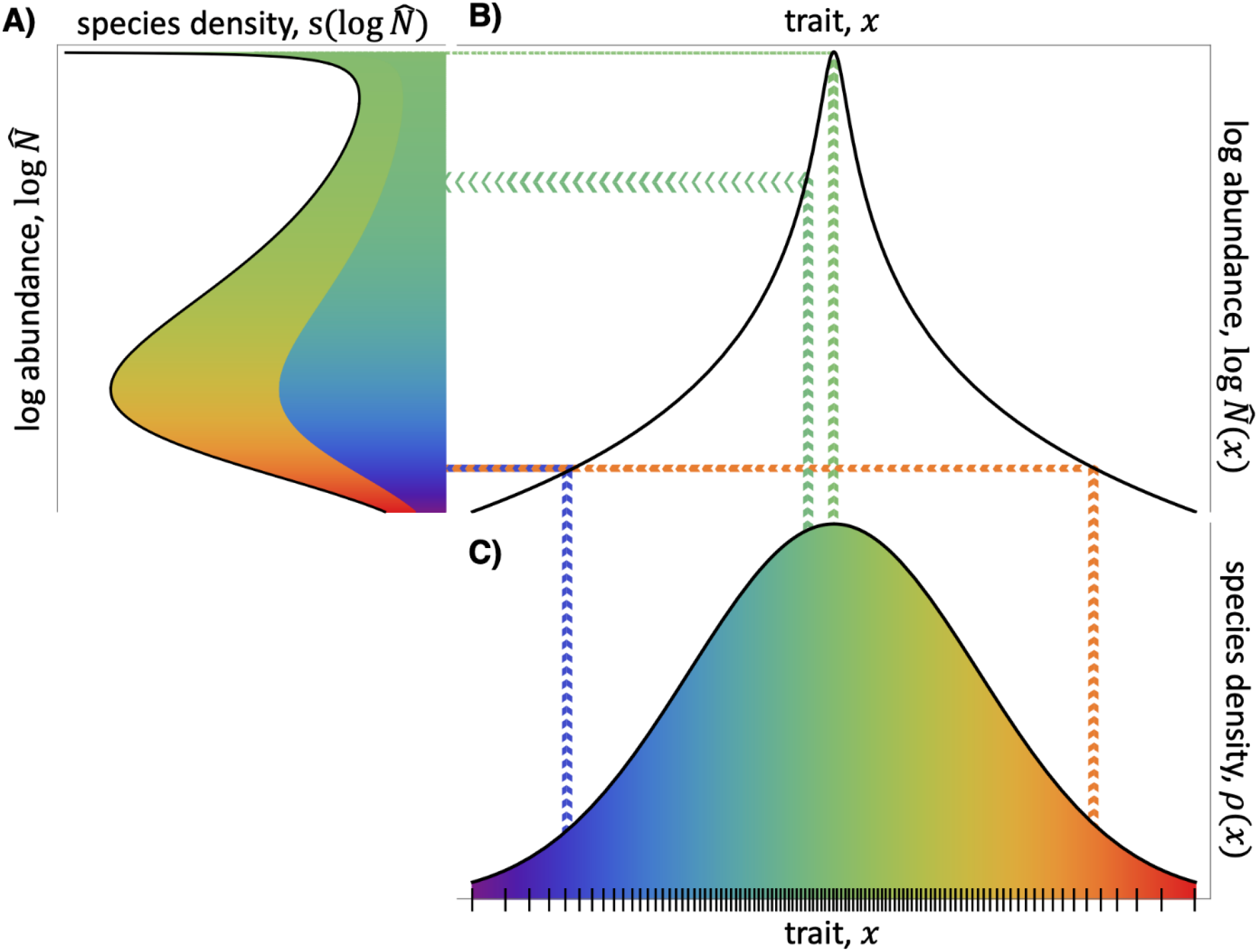
How the **A)** Preston-plot SAD results from the transformation by **B)** the TAD of **C)** the regional species-density function (tick marks indicate discrete species). Flat parts of the TAD concentrate species density in the SAD, whereas sloped parts diffuse it (indicated by the width of the beams reflected off the TAD).

Setting eqn. (4) equal to zero, the equilibrium TAD is formally given by

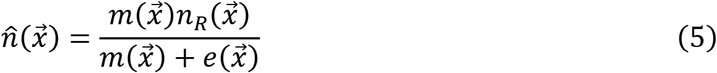

where 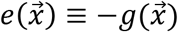 is the exclusion rate. Note that eqn. (5) is not a closed-form solution to eqn. (4), because the exclusion rate depends implicitly on the full TAD, 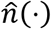. However, it can serve as the basis for analytical approximations, and highlights that abundance is determined by the balance of immigration and exclusion.

To proceed further, we need to specify the growth function *g*. We will begin with one of the simplest ecological interactions: competition in a single niche, with a quadratic intrinsic growth function of a scalar trait, *r*(*x*). This is an archetype of stabilizing selection (Lande & Arnold 1983), which is not only analytically tractable, but also useful as an approximation to more general situations. Defining *N*_tot_ = ∫ *n*(*x*)*dx* as the total abundance integrated over the entire trait axis, this results in

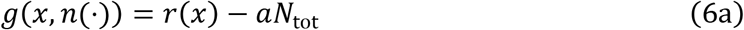

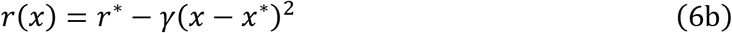

where *a* is the competition coefficient that scales population sizes, *r*\* is the maximal growth rate at the optimal trait value *x*\*, and *γ* quantifies the strength of stabilizing selection. In the absence of dispersal (indicated by zero subscripts), there is a single species (equivalently, a set of neutral species) with the optimal trait value *x* = *x*’ and total abundance 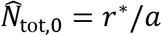, leading to a TAD consisting of a Dirac delta function: 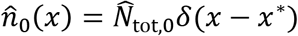.

Further assuming trait-independent dispersal *m*(*x*) = *m* and uniform regional abundance *n_R_*(*x*) = *n_R_*, we find that the equilibrium TAD is

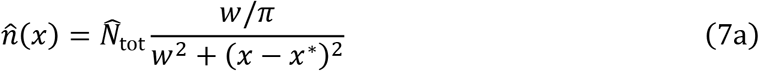

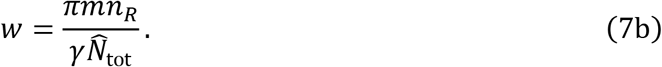

This is still not a closed-form solution, because the total abundance 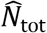 is not explicitly defined, but it shows that the TAD takes the form of a Lorentzian function, with characteristic width at half-maximum *w*. Lorentzians are bell-shaped curves superficially similar to Gaussians, but are heavy-tailed with a power-law exponent of −2 (Fig. 2A).

**Figure 2.**
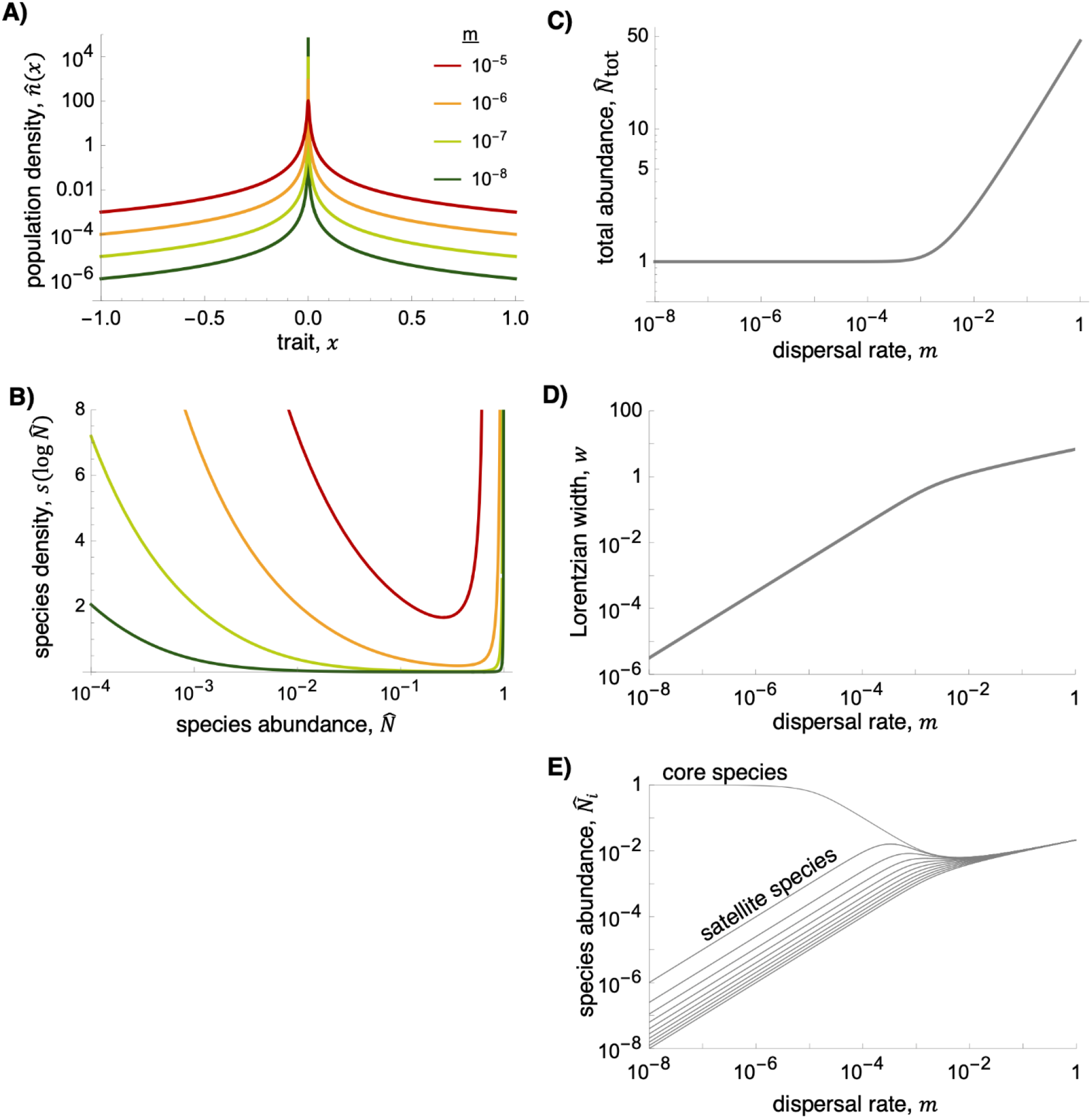
Our first example (one niche, quadratic fitness function, uniform regional TAD). Equilibrium **A)** TADs and **B)** SADs for four dispersal rates *m*. **C-E)** The effect of dispersal on **C)** total abundance 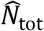, **D)** width of the Lorentzian TAD *w*, and **E)** abundance of 21 evenly spaced species between *x* = −1 and *x* = 1. Parameters: *r*\* = 1,*x*\* = 0,*γ* = 1,*n_R_*(*x*) = 100,*ρ*(*x*) = 100 so that *N_R_*(*x*) = 1.

In Appendix A we show that the total abundance 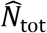 is the solution of a cubic equation, whose explicit form is non-insightful but is used for plotting results below. However, in the limit of low dispersal, it can be well-approximated as 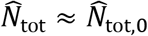, and in the limit of high dispersal as 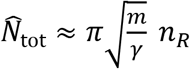, with a cross-over between these regimes around 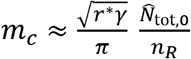 (Fig. 2C).

Note that total abundance diverges in the high dispersal limit in this simple example due to our assumption of uniform regional abundance (and therefore infinite regional abundance when integrated over the trait axis). Therefore, the low dispersal limit (*m* < *m_c_*) is more relevant here; other examples with finite regional abundance are well-behaved in both limits.

Figure 2 shows the effect of dispersal rate *m* on the equilibrium TAD 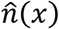 and SAD 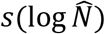, derived from the TAD following Box 1. The SAD has an increasing left-tail and a thin asymptote on the right, which represents the single core-species (Fig. 2B). As in the discrete-species case (eqn. 2a), the abundance of satellite species is proportional to dispersal rate (Fig. 2A,E). The width of the Lorentzian *w* increases with dispersal, as can also be seen from eqn. (7b) (Fig. 2D). Because the total abundance 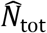 is approximately constant in the low dispersal regime (Fig. 2C), this implies that the density within the expanding core must decrease with dispersal (Fig. 2A). Therefore, for species with a given trait value close to the core species, density first increases linearly, then decreases as it enters the core community (Fig. 2E). In Appendix A we show that the core community can be defined as those within the characteristic width of the Lorentzian *w* of the peak.

#### Box 1: Converting TADs to SADs

Converting the TAD *n*(*x*) to an SAD requires two steps. First, we must bin individuals from the continuous distribution *n*(*x*) into species abundances *N*(*x*). Then log*N*(*x*) can be used to calculate a Preston-plot SAD, *s*(log*N*), which can be integrated to determine the number of species within a particular abundance class (Preston 1948, McGill et al. 2007). Both steps require the species-density function, *ρ*(*x*), which describes the density of species along the trait axis. Its inverse *δx*(*x*) = 1/*ρ*(*x*) gives the trait distance between species.

*<u>Step one</u>.* Except where *n*(*x*) is sharply peaked relative to the spacing between species, *δx*(*x*), *N*(*x*) can be well-approximated by the naïve approximation *N*(*x*) = *n*(*x*)/*ρ*(*x*). For the Lorentzian, this approximation is valid for core species when *w* » *δx*(*x*\*), and is always valid for satellite species (Fig. C1). Otherwise, we integrate *n*(*x*) across the distance between species:

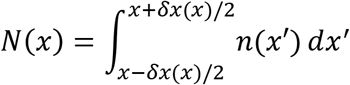

When *n*(*x*) is a Lorentzian distribution, this integral is 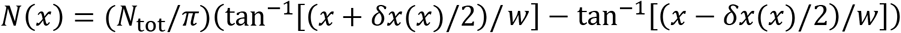.

*<u>Step two</u>.* The log-transformed TAD, log*N*(*x*), can be used to translate the species-density function, *ρ*(*x*), into a Preston plot SAD, *s*(log*N*). Mathematically, this is similar to transforming a random variable (Kobayashi et al. 2011). In general,

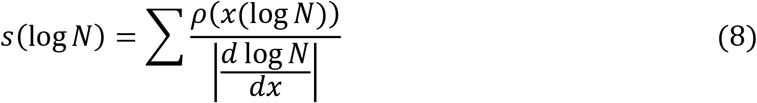

where the sum is over all branches of the inverse function *x*(log*N*), because the log-TAD is typically non-invertible. Note that flatter parts of the log-TAD result in more species with that abundance and sloping parts result in fewer, due to the denominator (Fig. 1).

In our example of a quadratic intrinsic fitness, one niche, trait-independent dispersal and uniform regional abundance and species density (*ρ*(*x*) = *ρ*) (Fig. 2), assuming *w* » *δx*(*x*\*), we find the SAD

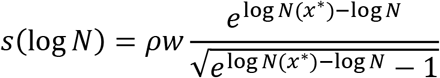

where 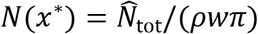 is the height of the Lorentzian TAD at its peak.

### Effect of regional species pool

The assumptions of uniform regional TAD *π_R_*(*x*) and species density *ρ*(*x*) were made above to facilitate analytical calculations, but clearly represent a special case. First, consider a nonuniform regional TAD. In this case, the regional TAD *π_R_*(*x*) undergoes a Lorentzian-like transformation by the exclusion rate e(x) (eqn. 5). Under the assumptions of one niche, a quadratic intrinsic fitness function (eqn. 6), and trait-independent dispersal, this results in the equilibrium TAD

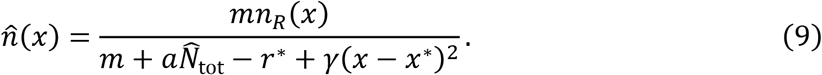

As before, this is not a closed-form solution due to the presence of the total abundance 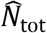. Although 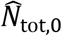 remains a good approximation at low dispersal rates, unfortunately, in general 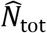 must be determined numerically. Note that in the absence of selection (neutrality, *γ* = 0), the local TAD is proportional to the regional TAD if dispersal is the same for all species.

Second, the technique given in Box 1 can be used to derive SADs from TADs in the case of nonuniform species density. As an example, consider a Gaussian species-density distribution, 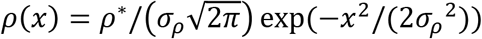, where the density of species decreases away from the origin (Fig. 3A). Assuming uniform regional species abundances *N_R_*(*x*) ≡ *N_R_*, the regional TAD is proportional to *ρ*(*x*): *n_R_*(*x*) = *N_R_ρ*(*x*). In this case the TAD has a narrow spike at the optimal trait value (Fig. 3B); it superficially resembles that of our previous example (Fig. 2A), but decreases more sharply due to the lower regional abundance away from *x* = 0. However, in contrast to the case of uniform regional species density, the resulting SAD is now unimodal aside from the singular core species at the right (Fig. 3C), due to the low density of species in the tails of the species-density distribution *ρ*(*x*).

**Figure 3.**
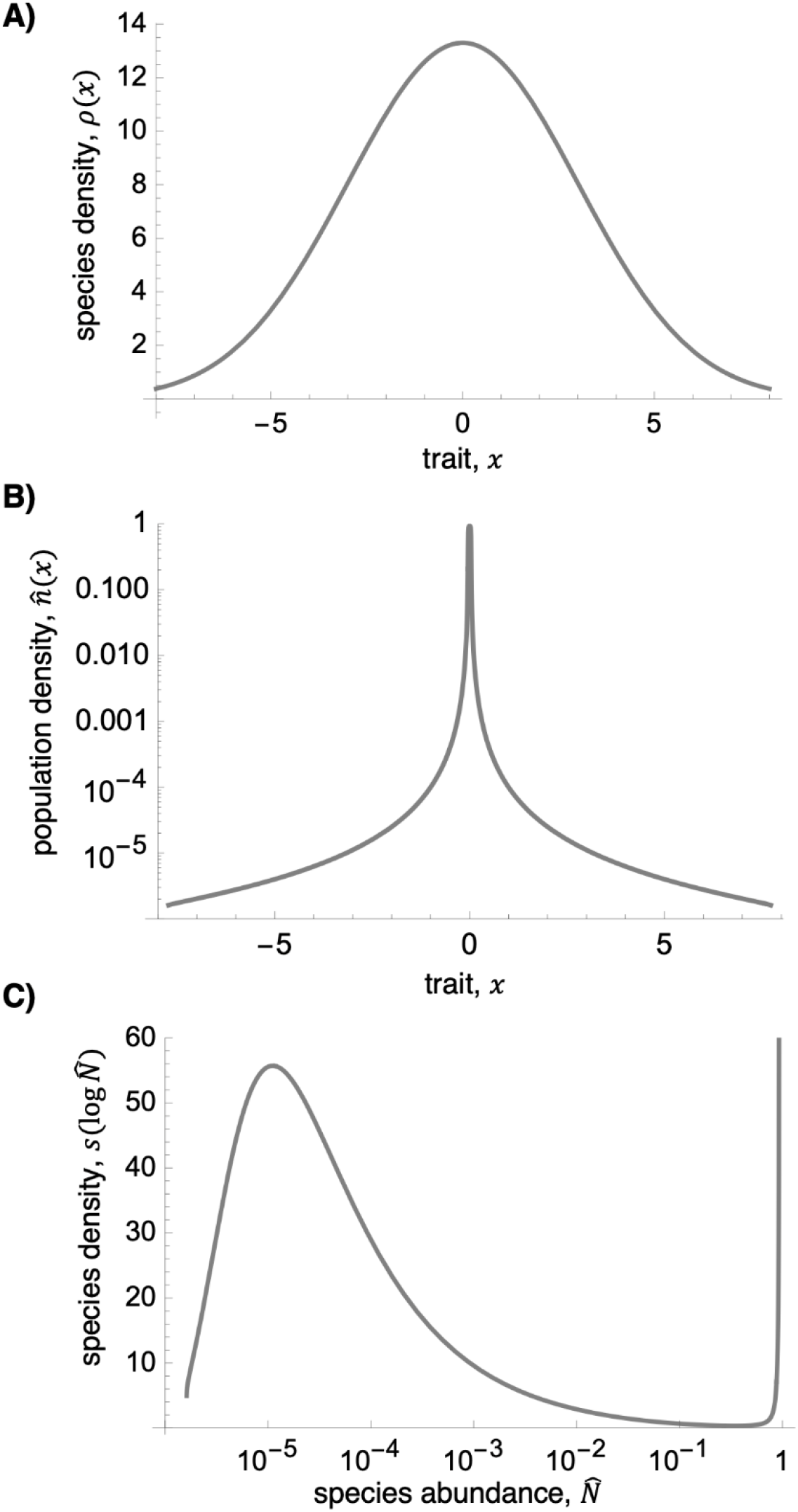
Our second example (one niche, quadratic fitness function, Gaussian regional TAD). **A)** Regional TAD, **B)** local TAD, and C) unimodal SAD. Parameters as in Fig. 2 except *σ_ρ_* = 3,*ρ*\* = 100, *N_R_* = 1, *m* = 10^−4^.

### Other interaction models (non-quadratic r, multiple niches)

The previous examples all assume one niche and a quadratic intrinsic fitness function, representing an analytically tractable model of stabilizing selection. However, our general traitbased framework of eqn. (4) is applicable to a much wider range of ecological scenarios. Here we give two more examples to show how the details of local interactions affect TADs and SADs. Although no analytical results are available in general, in Appendix C, we show that these TADs can be approximated as Lorentzians near a peak.

First, consider one niche but a Gaussian (non-quadratic) intrinsic fitness function, 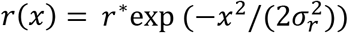. This can be applicable if the trait affects fitness only through the birth rate, because birth rates must be non-negative, so the fitness function must be bounded from below. Figure 4A shows that the equilibrium TAD is peaked around the optimum *x* = 0 but is also bounded from below. This can be understood from the exclusion rate at equilibrium (Fig. 4B), which approaches a constant maximum value, no matter how maladapted a species is. This flat range of the TAD translates into a vertical asymptote at 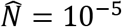 on the left side of the SAD (Fig. 4C).

**Figure 4.**
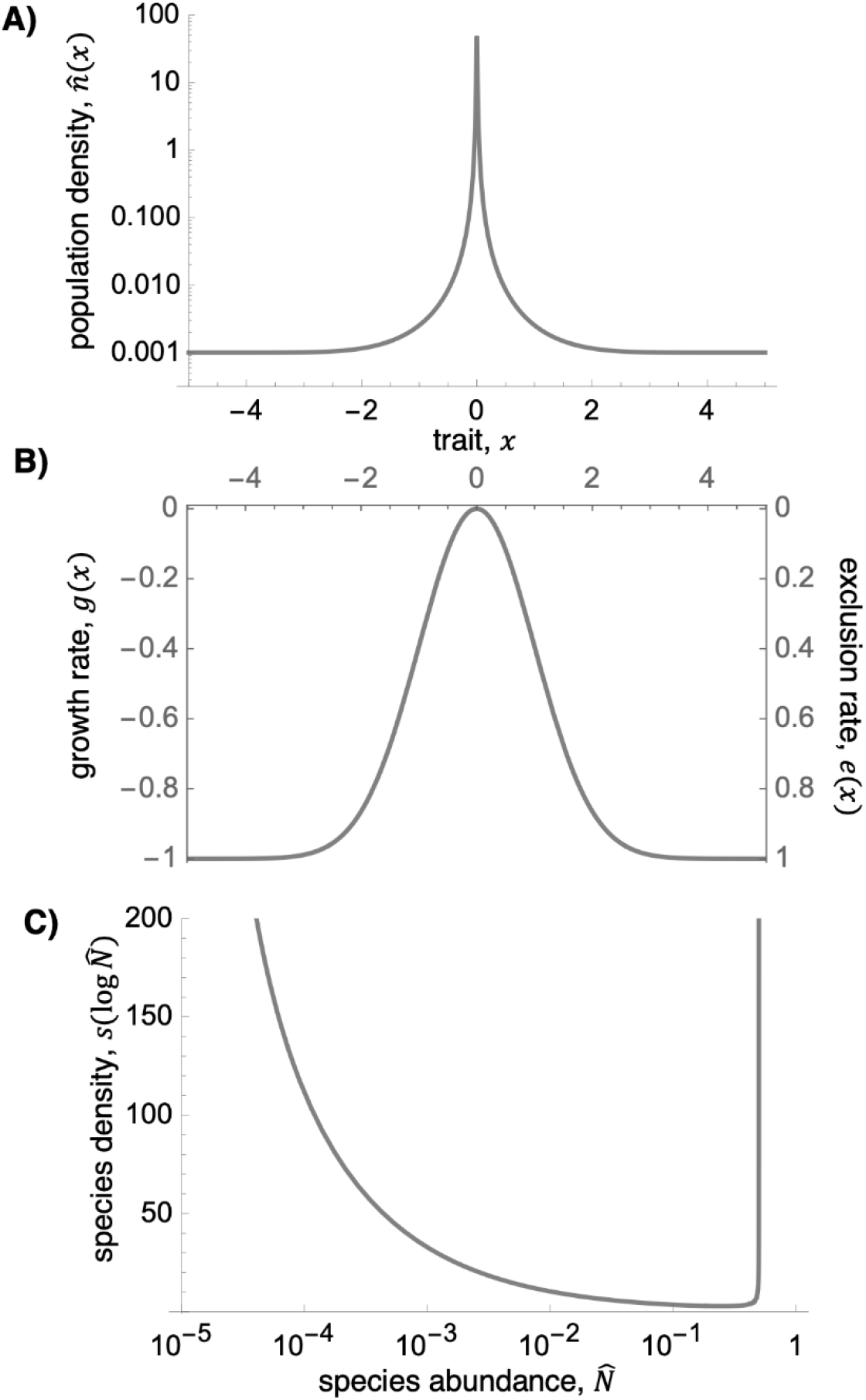
Our third example (one niche, Gaussian fitness function, uniform regional abundance). **A)** TAD, **B)** growth/exclusion rate, **C)** SAD. Parameters as in Fig. 2 except *σ_r_* = 1,*m* = 10^−5^.

Finally, we relax the assumption that there is only one niche by letting the strength of competition *a* decline with the difference in traits between interacting species. That is,

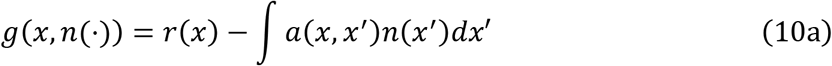

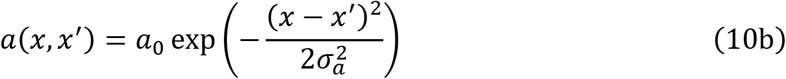

This trait-based Lotka-Volterra competition model is a classic model for niche differentiation and has been extensively explored (Scheffer & van Nes 2006, Ranjan & Klausmeier 2022). In the absence of dispersal, when 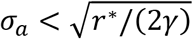 there is an evolutionarily stable community (ESC) consisting of more than one species (Ranjan & Klausmeier 2022). In our example, the ESC consists of three evenly-spaced core species that sit at local invasion fitness maxima (Fig. 5A).

**Figure 5.**
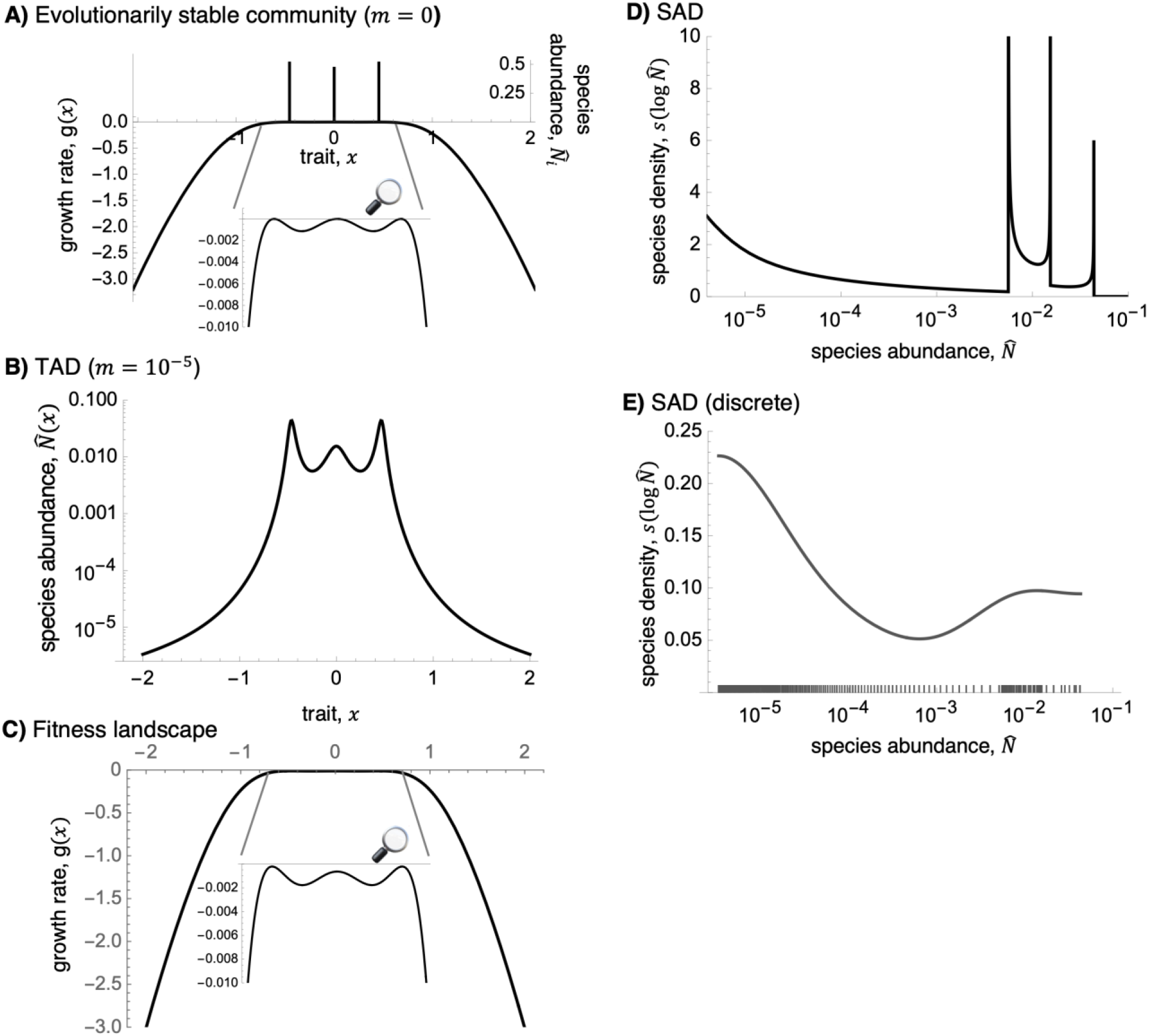
Our fourth example (localized competition, quadratic fitness function, uniform regional abundance). **A)** The evolutionarily stable community reached in the absence of dispersal and the corresponding invasion-fitness landscape. **B)** TAD with dispersal and **C)** the corresponding fitness landscape. **D)** Continuum-based SAD. **E)** Discrete-species-based SAD.

Because the continuum model eqn. (4) would need to be discretized before solving numerically, we directly simulate eqn. (3) with a discrete set of species. With small dispersal, the TAD has three locally Lorentzian peaks (Fig. 5B), whose heights and widths are largely determined by the non-dispersal abundance and curvature of the fitness function (Appendix C). Overall, the TAD is determined by eqn. (5), with the exclusion rate playing the central role given our assumption of uniform regional abundance and species density (Fig. 5C). Because the exclusion rate for species between the three core species is very small (on the order of 10^−3^), it takes only a small immigration rate to boost these nearly neutral species to appreciable densities (Fig. 5B).

Like the TAD, the SAD is multimodal (Fig. 5D): the two rightmost peaks in the SAD correspond to the peaks in the TAD (two of which have the same height), whereas the third peak from the right in the SAD corresponds to the local minima of the TAD. However, this fine detail may be difficult to detect in empirical SADs. Fig. 5E shows a SAD fitted using kernel density estimation on a discrete set of 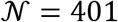 species, indicated by ticks on the *x*-axis. The narrow peaks of the theoretical SAD (Fig. 5D) cannot be distinguished, and merge into one wide peak.

### Controls on the distribution of satellite species

Because most species in local communities are rare (McGill et al. 2007), to understand the bulk of TADs and SADs we can focus on the satellite species, which greatly simplifies the following analysis. Let *e*_0_(*x*) ≡ −*g*_0_(*x*) denote the exclusion rate (negative growth rate) in the absence of dispersal. This is determined by the invasion fitness function used to find an evolutionarily stable community in adaptive dynamics (Geritz et al. 1998, Klausmeier et al. 2020). For satellite species with *e*_0_(*x*) » *m*, the rate of exclusion is great relative to the dispersal rate, so we can simplify eqn. (6) to the zeroth-order approximation analogous to eqn. (2a)

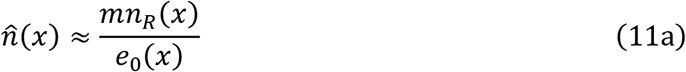

or equivalently,

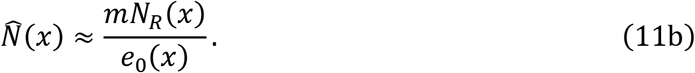

In contrast to the implicit eqn. (5), eqn. (11) gives explicit formulas for the TAD of the weakly excluded satellite species. Therefore, it can give analytical insights into the relative abundance of rare species in a wide range of scenarios, where the exact formula cannot be solved. Fig. C1 compares the approximation to the TAD given by eqn. (11) and the resultant SAD, to the exact results given by eqn. (5) under naïve and integral transformations (Box 1). It is an excellent approximation, except where eqn. (11) diverges due to dividing by *e*_0_(*x*\*) = 0 at the core species.

Of particular historical interest is the behavior of the left tail of the SAD, which describes the relative abundance of rare species (Fisher et al. 1943, Preston 1948). To determine whether the left tail increases without bound, approaches zero (implying an at-least-unimodal SAD), or approaches a non-zero constant (as the log-series SAD does), we examine the limit of eqn. (8) using eqn. (11), as *x* → ±∞. If *N*(*x*) → *N*_min_ > 0, as occurs when *e*_0_(*x*) is bounded from above, then the SAD has a vertical asymptote at log *N*_min_ (Fig. 4).

Otherwise, if *N*(*x*) → 0 as *x* → ±∞, we assume that the asymptotic behavior of each function of *x* can be described by a power function times a generalized exponential function; that is,

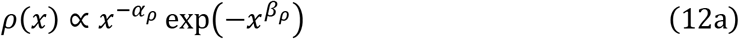

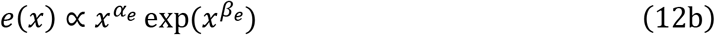

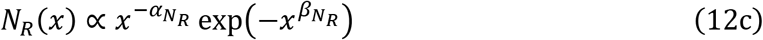

with all exponents non-negative (*α* ≥ 0 and *β* ≥ 0). We find (Appendix D), that if the speciesdensity function *ρ*(*x*) decays faster than a power law (*β_ρ_* > 0), then *s*(log*N*) → 0 as *N* → 0, as the Gaussian *ρ*(*x*) does in Fig. 3C. If the species-density function *ρ*(*x*) decays more slowly, as a power law (*β_ρ_* = 0), then the left tail of the SAD depends on exponent of the species-density function relative to the larger exponent of the regional population-density function and the exclusion function:

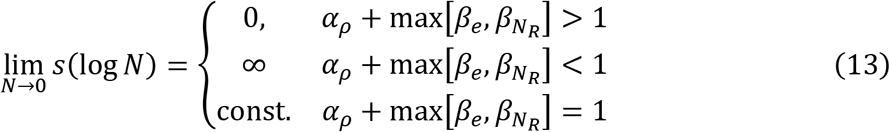

A special case: if *ρ*(*x*) and *N_R_*(*x*) are constant 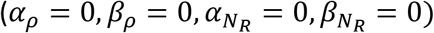, then *s*(log*N*) → 0 if the exclusion function *e*_0_(*x*) increases faster than exponentially (*β_e_* > 1) and *s*(log*N*) → ∞ if it declines slower than exponentially (*β_e_* > 1) (as our example in Fig. 2B).

## Discussion

What determines the shape of species abundance distributions (SADs)? Our analysis suggests that the simple answer is “there is no simple answer.” Both local processes (environmental filtering and species interactions) and the properties of the regional species pool (species abundances and traits) jointly determine the distribution of commonness and rarity in a community. These factors span from local to landscape and biogeographic scales, each subject to tremendous variation and historical idiosyncrasy. Therefore, it seems highly unlikely that there is a universal, parameter-sparse SAD that is applicable across all communities. Despite the manifold variety of possible SADs it can produce, our framework identifies key factors that shape them — rates of exclusion and dispersal, regional population abundance, and the species-density distribution. Moreover, it makes a number of empirically testable predictions outlined below.

First, although our framework can produce a multitude of SAD shapes, some general predictions exist. It predicts that at low dispersal rates, the SAD will have a gap between the most abundant core species and the rare species that make up the bulk of the community. It also can explain multimodal SADs (Dornelas & Connolly 2008, Antão et al. 2017), generalizing the findings of Vergnon et al. (2012) and Rael et al. (2018). If the scaling exponents of the exclusion function and the regional species pool are known, it predicts the behavior of the lefthand tail of the SAD (rare species) (eqn. 13).

Second, it predicts that at low immigration rates, the abundance of rare, satellite species will vary linearly with the immigration rate (eqn. 2), while the abundance of common, core species will be largely unaffected (Fig. 2E). An important corollary is that satellite species cannot persist in the absence of immigration. The immigration rate is the product of the dispersal rate *m_i_* and the regional abundance *N_R,i_*. High dispersal leads the local community to converge with the regional metacommunity. These predictions could be tested by manipulating the immigration rate experimentally. Patrick (1967) showed that increased immigration led to SADs with more rare species of stream diatoms. Further experimental tests in different systems are needed.

Third, our general framework predicts that the identity and abundance of dominant core species will be determined by their traits and environmental conditions. In contrast to neutral theory, replicate local communities with similar environmental conditions should have the same dominant species. The most informative observational evidence would come from cases where the local communities deviate strongly from the regional metacommunity, since neutral theory predicts that regionally abundant species will also tend to be locally abundant. Experimental manipulation of environmental conditions or starting new communities would provide an even more powerful test of this prediction. Murray et al. (1999) found that 91-95% of locally rare plant species were locally abundant at another site within their range, consistent with our theory.

Fourth, it predicts that the exclusion rate (negative invasion/growth rate) strongly affects species abundances. Invasion rate is a central concept in ecological theory and can be experimentally measured in a number of ways (Grainger et al. 2019), providing another approach to test the general framework.

More-specific models can generate more-specific hypotheses. Because our general framework can accommodate a wide variety of particular models through the growth function *g*, predictions of how TADs and SADs vary along environmental gradients, such as nutrient and stress gradients, can be made. Thus, verbal hypotheses for how SADs should vary with pollution (Gray 1979) could be made more precise, leveraging the broad base of trait-based ecological theory (Klausmeier et al. 2020).

Population-dynamic theoretical explanations of SADs invoke one (or both) of two alternative sources of variation in species abundance: *chance* and *fate.* In chance-based explanations, species have identical parameters (symmetric models, which include non-interacting species and neutral models as special cases) but include either environmental (Engen & Lande 1996ab) or demographic stochasticity (Kendall 1948, Caswell 1976, Hubbell 2001). In fate-based explanations, dynamics are deterministic, but species have different parameters, either randomly assigned (Wilson et al. 2003, Wilson & Lundberg 2004, Zhou & Zhang 2008) or determined by traits (Vergnon et al. 2012, our approach). A few models combine both sources of variation (Hughes 1986, Haegeman & Loreau 2011, Rael et al. 2018). While both approaches can produce realistic SADs, an important distinction concerns the role of species identity. In chance-based explanations, all species will eventually experience the entire range of abundances in the SAD, whereas in fate-based explanations, species will be consistently common or rare in a particular environment.

Of Vellend’s four fundamental processes that shape ecological communities (2010), our approach is based on selection and dispersal. Because we focus on local community dynamics on ecological timescales, we do not explicitly model speciation, but it can be considered implicitly as a determinant of the regional species pool.

Selection embodies diverse interspecific interactions with a seemingly endless number of variations, from abstract, general models like Lotka-Volterra, to those tailored to particular ecosystems. Indeed, the majority of ecological theory includes only selection, due to its inherent richness. Although our examples are based on Lotka-Volterra competition (with one or multiple niches), our analytical framework (in general, eqn. 1; assuming trait-based interactions, eqn. 3) can be applied to a much wider range of ecological scenarios though the growth function *g*.

We model dispersal as the exchange of individuals with the fixed regional metacommunity. In particular, immigration plays the central role in shaping local community structure through mass effects (Shmida & Wilson 1985). While some studies invoke immigration to explain the distribution of transient species in SADs (Magurran & Henderson 2003), previous theory has shown that even a small amount of immigration can translate into large population sizes in sink populations (Pulliam 1988, Holt 1993, Gonzalez & Holt 2002). This inflationary effect can be seen in previous models of SADs (Kendall 1948, Hughes 1986, Loreau & Mouquet 1999, Gravel et al. 2006, Scheffer & van Nes 2006) and forms the basis of our analytical theory (eqn. 2). Note that while we consider spatial mass effects through dispersal, temporal mass effects in the form of a seed bank of resting stages (Lennon & Jones 2011, Alexander et al. 2012) would act similarly.

Unlike neutral theories (Caswell 1976, Hubbell 2001), we neglect ecological drift (demographic stochasticity). The primary reason is to contrast fate-based explanations of SADs with chancebased explanations. Demographic stochasticity is most important in small populations, so our approach is most suitable for communities with high total abundance, but may need to consider drift in the rarest species. Future research that integrates all three fundamental ecological processes (selection, dispersal, and ecological drift) can build off the analytical foundations of our current framework. In particular, rare satellite species that do not affect others can be modeled as independent birth-death-immigration processes that lead to negative-binomially distributed abundances (Kendall 1948). Because we focus on the theoretical processes underlying SADs, we also neglect random variation due to the sampling process, which will be required for rigorous comparison with empirical data (Bulmer 1974). We anticipate the effect of limited sampling will be qualitatively similar to the unveiling of the lognormal distribution noted by Preston (1948).

The fitness (growth) function is a central concept in eco-evolutionary frameworks such as adaptive dynamics (Geritz et al. 1998). However, the fitness function is often used only as a means towards the end of determining the evolutionarily stable community (ESC) (Brown & Vincent 1987, Klausmeier et al. 2020). With immigration from the regional metacommunity, the ESC represents the bones of the community (core species), while the fitness function helps shape the flesh (satellite species). Rather than being ultimately discarded as in adaptive dynamics, the fitness function itself is a key result that gives valuable information on the bulk of the community in the presence of immigration.

Our framework connects SADs with species- and trait-based community models, setting the stage for future theoretical extensions, including incorporating temporally varying environments (random: Chesson 2000, seasonal: Kremer & Klausmeier 2017) and structured populations (discrete: Caswell 2001, continuous: De Roos & Persson 2001) into SAD theory. While we have focused on long-term equilibrium behavior, models such as eqn. 1 and 3 can also be used to study transient dynamics (DeAngelis & Waterhouse 1987, Hastings 2004). Thus, our framework lays a general theoretical foundation for the study of SADs, allowing them to become fully integrated into the mainstream of theoretical ecology.

## Acknowledgments

We thank Jonas Wickman for helpful comments on the manuscript. This research was supported by the Simons Foundation grant 343149, NSF grant DEB 17-54250 and NASA grant 80NSSC18K1084.

## Appendix A: Mathematical details of example 1

In this appendix we provide details of the derivation of our results in the case of one niche, quadratic fitness function, and uniform regional abundance. Following the approach in the main text, we have the following expression for the equilibrium TAD:

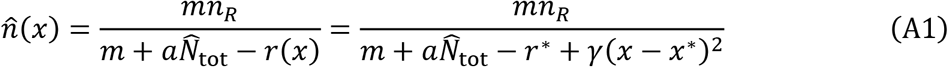

where

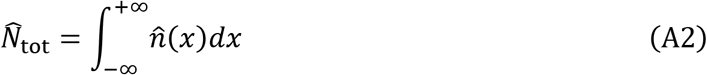

Integrating eqn. (A1) over *x* yields an implicit equation for the total density 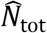

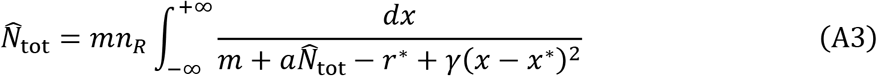

which can be rewritten as

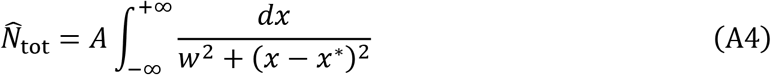

with width

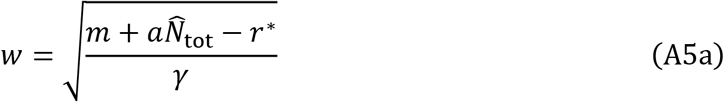

and amplitude

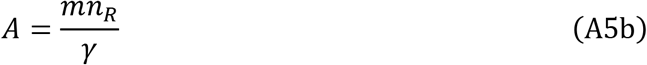

We recognize the primitive of the inverse tangent function in the integrand of eqn. (A4), which can be integrated to

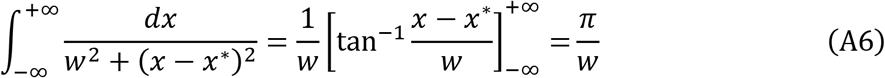

This leads to an expression relating the width and amplitude of the Lorentzian distribution with the total abundance:

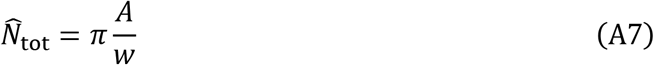

We substitute eqns. (A5) and note that *w* itself is a function of 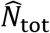 (eqn. A5a), so squaring this expression and rearranging terms, we get

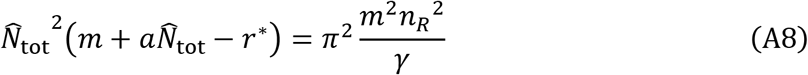

Now, let us use 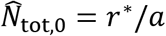 to rewrite this equation as

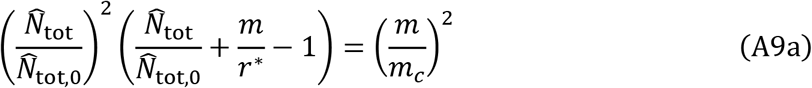

with

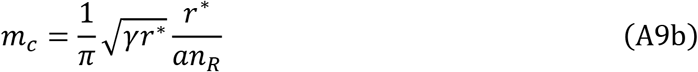

The implicit equation (A9) does not admit a particularly insightful explicit solution. However, we see that there are two natural scales for *m* to influence 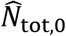: the first one, on the left-hand side of eqn. (A9a) compares *m* to *r*\* to see when emigration has a significant effect. The second one, on the right-hand side, compares *m* to *m_c_* to see when immigration has a significant effect on total abundance.

To understand the behavior for small *m*, let us Taylor expand the implicit equation (A9) to the second order in *m,* using

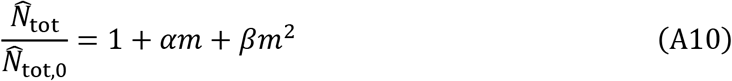

with the coefficients *α* and *β* to be determined. We get:

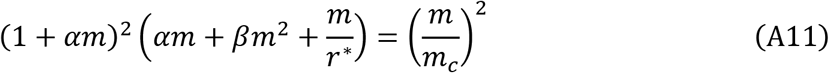

which reduces to:

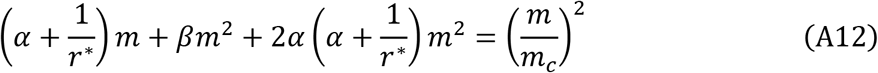

from which we deduce that *α* = –1/*r*\* and *β* =1/*m_c_*^2^. To conclude:

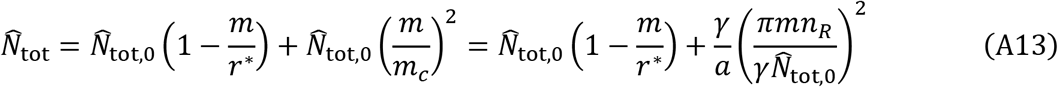

When eqn. (A13) is substituted back into eqn. (A5a), it gives us the lowest order approximation for the width *w*:

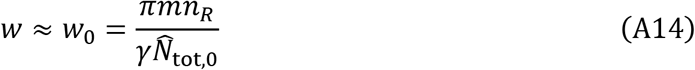

When eqn. (A14) is substituted back into the TAD (eqn. A1), we get at the lowest order:

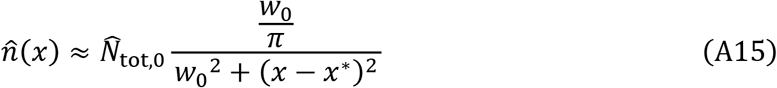

which is eqn. (7) in the main text with 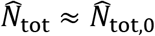 and *W*≈*W*_0_.

Let us now look at the response of the TAD (eqn. (A15)) to increased dispersal in this small dispersal limit. A small increase in dispersal widens the trait abundance distribution 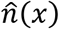 proportionally to *m*, due to the dependence of *w*_0_ on *m* (eqn. A14). At the same time, total abundance remains approximately constant at 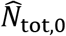, so that the amplitude of the distribution *A* must decrease with 1/*m*. Interestingly, this implies that the abundance of the strategies close to the optimal strategy *x*\* decrease as the abundance of satellite species increases. This is quantified by computing the response of the abundance of a strategy with trait *x* to increased dispersal *m*, formally given by

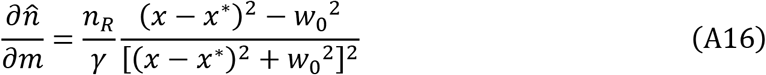

As we can see from eqn. (A16), strategies with trait x within the characteristic width distance from the optimum *x*\*, i.e. such that |*x* — *x*\*| < *w*_0_, see their abundance decrease with increasing immigration. Conversely, strategies with trait *x* outside of the characteristic width distance from the optimum *x*\*, i.e. such that |*x* – *x*\*| > *w*_0_, see their abundance increase. This justifies the operational definition of core and satellite species provided in the main text.

Conversely, in the large dispersal limit, eqn. (A9) simplifies to

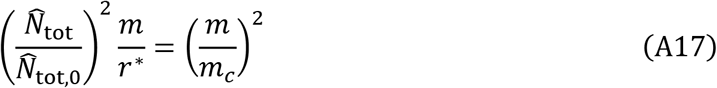

so that

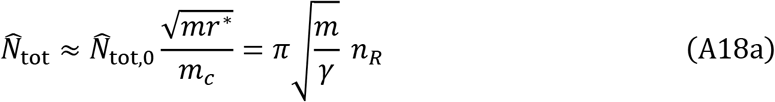

and

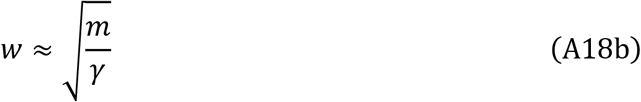

as seen in Fig. 3A&B.

## Appendix B: Approximating peaks as Lorentzians

In this appendix we present the calculation that leads to the TAD for a general Lotka-Volterra system in the small dispersal limit. Our starting point is the trait-based Lotka-Volterra model

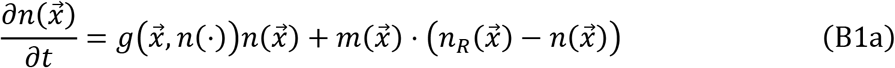

with growth function

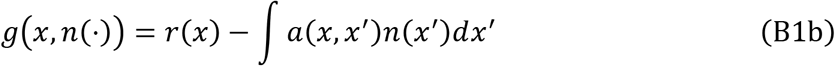

and competition kernel

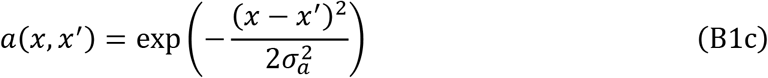

We know that in the absence of dispersal (*m* = 0), the equilibrium solution 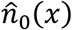 of the system above is a community made of a (potentially infinite) set of isolated peaks at locations *x*_0,*i*_ with densities 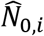 (see Fig. 5A for an example), taking the form

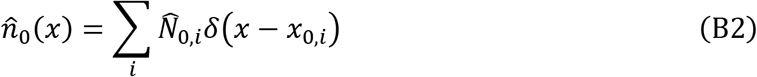

In the presence of immigration, the trait abundance distribution 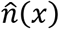 is formally given by:

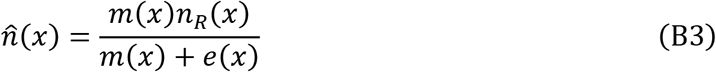

where the exclusion function *e*(*x*) implicitly depends on 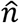:

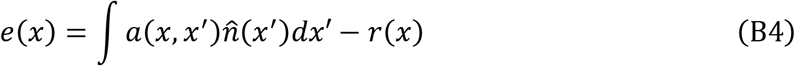

As we explained in the main text, there is no closed form solution for 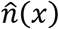. Indeed, solving explicitly for 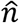 means solving a non-linear integral equation, which in general cannot be done analytically. Moreover, interactions in the multiple niche case are not all funneled through a single number 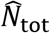 like in the single niche case, which prevents us from reducing the dimensionality of the problem. Still, let us rewrite eqn. (B3):

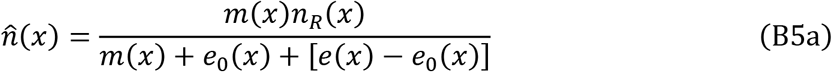

where

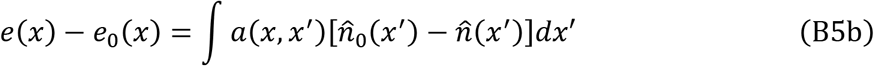

with 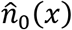 the solution of the continuum model in the absence of dispersal introduced above.

This gives:

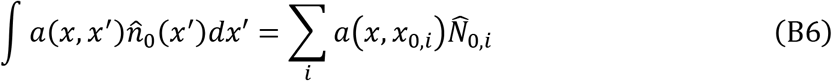

With the results on the one niche, quadratic fitness function, and uniform regional abundance case in mind (Eqn. (7) and Appendix A), we expect the quantity *e*(*x*) – *e*_0_(*x*) to stay small when dispersal *m* is small. For this reason, we assume that the main effect of dispersal on the exclusion function is to change the total abundance 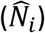 and location (*x_i_*) of each peak by small amounts 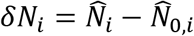 and *δx_i_*, = *x_i_* – *x*_0,*i*_. Thus the remaining goal of this analysis is to solve for *δN_i_* and *δx_i_*.

Substituting into (B6), we find

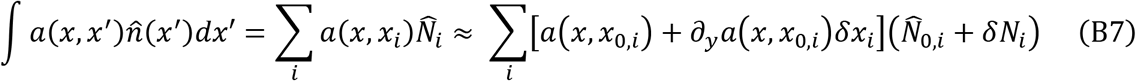

using the notation

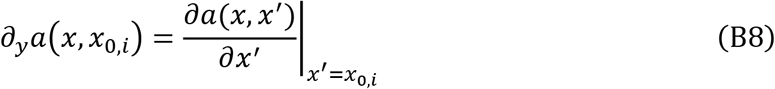

where the *∂_y_* stands for the derivative of *a* with respect to its second argument. After keeping only the first-order terms, we get

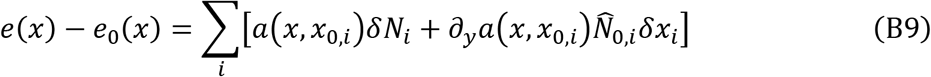

When substituted in the TAD, this gives

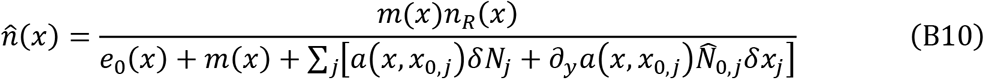

This equation turns into an implicit equation on *δN_i_* and *δx_i_* after convolution of both sides with the competition kernel *α*(*x, x*’):

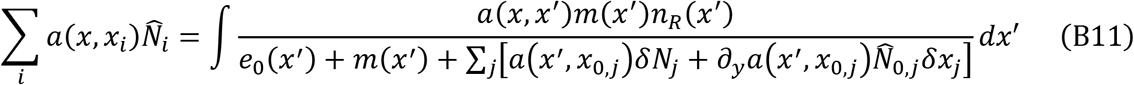

Note that eqn. (B11) has to be true for any *x*. The right-hand side of this self-consistent equation cannot be integrated directly. However, the small dispersal limit allows for further simplifications. Indeed, we expect the integrand of the right-hand side of eqn. (B11) to only be significantly different from zero in the small neighborhood of the *x*_*i*,0_, because the TAD should be strongly peaked around the location of the species of the ESC when dispersal is small. This means that this integral can be decomposed as a sum of integrals around the neighborhoods of the *x*_*i*,0_. In addition, *e*_0_(*x*) can be Taylor expanded around each *x*_*i*,0_ into:

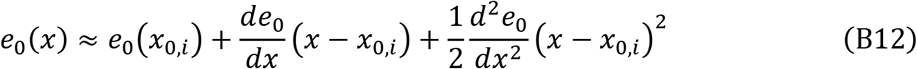

However, we know that the *x*_0,*i*_ are both the zeros and the maxima of *e*_0_(*x*), so eqn. (B12) simplifies to

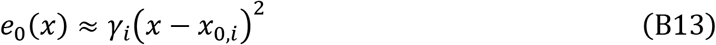

where the notation 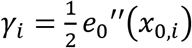, the local curvature of the fitness landscape, is used to strengthen the analogy with the quadratic *r* case used in eqn. (6b) and Appendix A. When eqn. (B13) is substituted into eqn. (B11), we obtain:

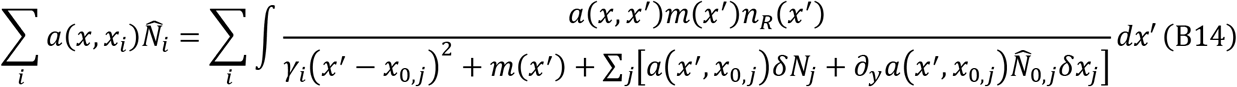

Similarly, we can Taylor expand the numerator of the integrand (using the unconventional approach of putting these developed terms in the denominator, to facilitate their integration with the rest of the denominator of the integrand):

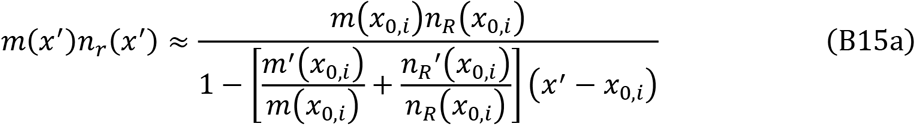

and

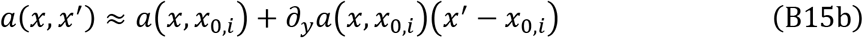

Similarly, let us Taylor expand the other terms in the denominator of the integrand and only keep terms of at-most order two in *m* and *x*’ – *x*_0,*i*_:

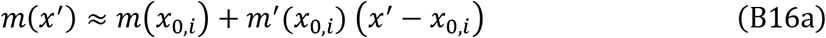

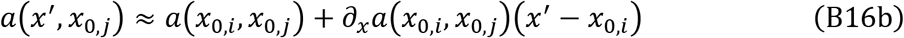

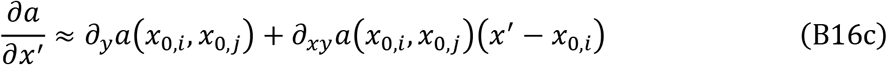

so that eqn. (B14) can be rewritten as

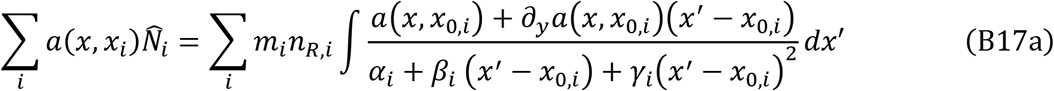

with the newly introduced coefficients

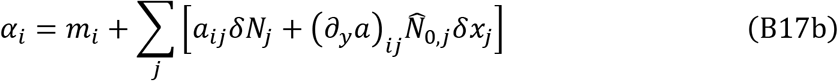

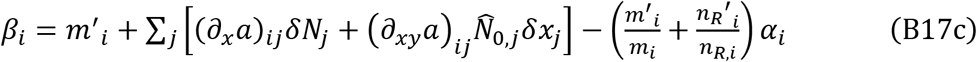

After introducing the simplifying notation 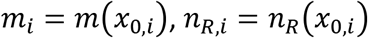, and 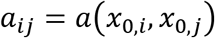 and the similar notation for their derivatives, eqn. (B17a) factors into

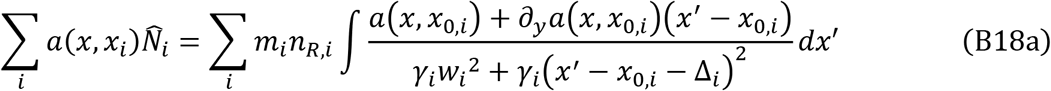

with the offset

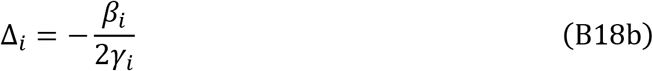

and the width

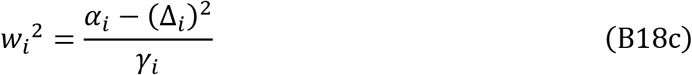

Now, using the Taylor expansion of eqn. (B7), the left-hand side of eqn. (B18a) can be rewritten as

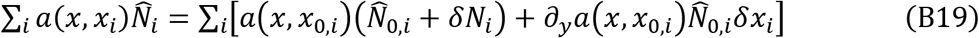

From there, eqn. (B18a) tells us that a sum of functions on the left-hand side (eqn. (B19)) is equal to another sum of function on the right-hand side. Assuming that these functions *a*(*x,x*_0,*i*_) and *∂_y_a*(*x,x*_0,*i*_) form independent bases, we can identify their coefficients by setting them equal to each other term-by-term, i.e., for the coefficients in front of the *a*(*x,x*_0,*i*_), we get

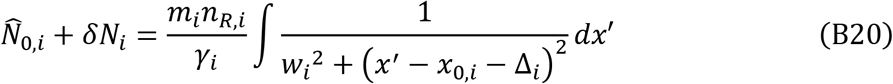

and for the coefficients in front of the *∂_y_a*(*x, x*_*i*, 0_), we get

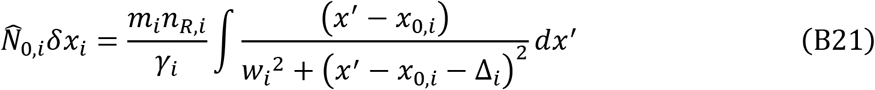

We recognize a Lorentzian function in each integrand in eqn. (B20) and eqn. (B21), which can be integrated in eqn. (B20) to give

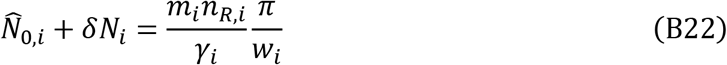

At the lowest order, this gives the width of the Lorentzian as:

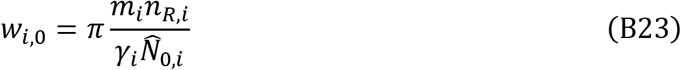

Notice how this expression for the width of the TAD around the peak of species *i* is formally identical to the result of eqn. (7b) obtained for the quadratic growth function case, justifying the claims in the main text, with the nuance that *m_i_, n_R,i_*, and *γ_i_* are, respectively, the dispersal rate, the regional population-density and the local curvature of the fitness landscape, all evaluated at *x*_0,*i*_.

Similarly, eqn. (B21) can be rewritten as

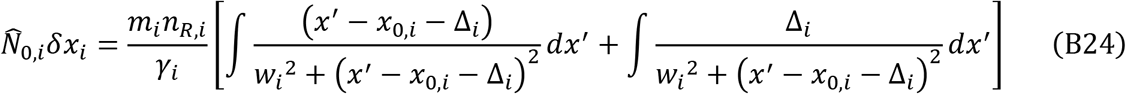

The first integral in eqn. (B24) is, as the integral of an odd function, equal to zero, while the second integral is similar to the one in eqn. (B20), giving after integration

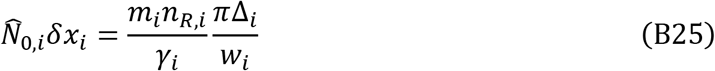

Using the lowest order approximation given by eqn. (B23), eqn. (B25) simplifies to Δ*i* = *δx_i_*,, which is quite intuitive: the shift *δx_i_*, in the competition kernel of the exclusion function has to coincide with the shift Δ_*i*_, that emerges in the denominator of the Lorentzians in eqns (B20) and (B21). Going back to 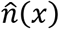 in eqn. (B5) and Taylor expanding at the neighborhood of a *x*_0,*i*_, we get at the lowest order

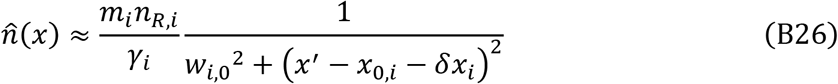

This is enough to conclude that the TAD for a general Lotka-Volterra model under small dispersal is a combination of localized Lorentzians with shifted peaks at *x*_0,*i*_ + *δx_i_*, and an effective stabilizing selection 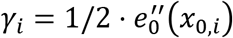 corresponding to the curvature of the fitness landscape at *x_oi_*.

Note that integrating each local Lorentzian in eqn. (B26) leads to a total abundance in peak *i* of 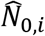. That is, it gives only the zeroth order approximation, which doesn’t help solving for the first order change *δN_i_*, due to small dispersal. Importantly, also notice that the value taken by *δx_i_*, has not been solved for yet.

Therefore, to solve for *δN_i_*, and *δx_i_*, we have to get back to eqns. (B18b) and (B18c) uncovered earlier (substituting Δ_*i*_ with *δx_i_*):

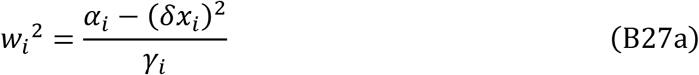

and

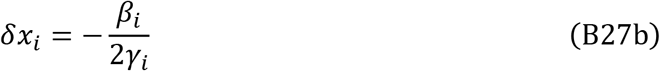

We expect both *δN_i_*, and *δx_i_* to have non-zero first order terms in *m*, which means that eqn. (B27a) directly gives *α_i_* = 0 to the first order (the two other terms, because they are squares, are second order), leading to the first set of linear constraints linking *δN_i_*, and *δx_i_*

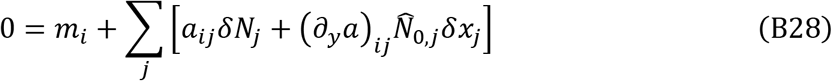

In addition, using the identity *α_i_* = 0 to the first order in eqn. (B27b) gives us a second set of linear constraints linking *δN_i_*, and *δx_i_*

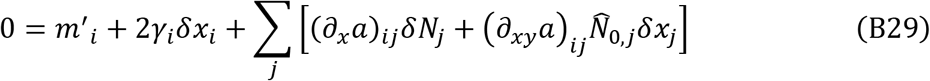

To conclude, *δN_i_*, and *δx_i_* satisfy a system of coupled linear equations given by eqns. (B28) and (B29), which can be inverted numerically to give the values of the peak displacements and the change in total density associated with each peak.

Importantly, these first order responses of 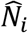, and *x_i_* to small dispersal are a consequence of emigration directly affecting growth, leading to a reorganization of interactions between the peaks, not immigration. Indeed, we believe that a small perturbation −*m*(*x*) to the growth rate of the original trait-based model *in absence* of immigration would lead to the exact same response of 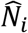 and *x_i_*.

## Appendix C: Comparing the zeroth-order approximation to the actual solution

**Figure C1.**
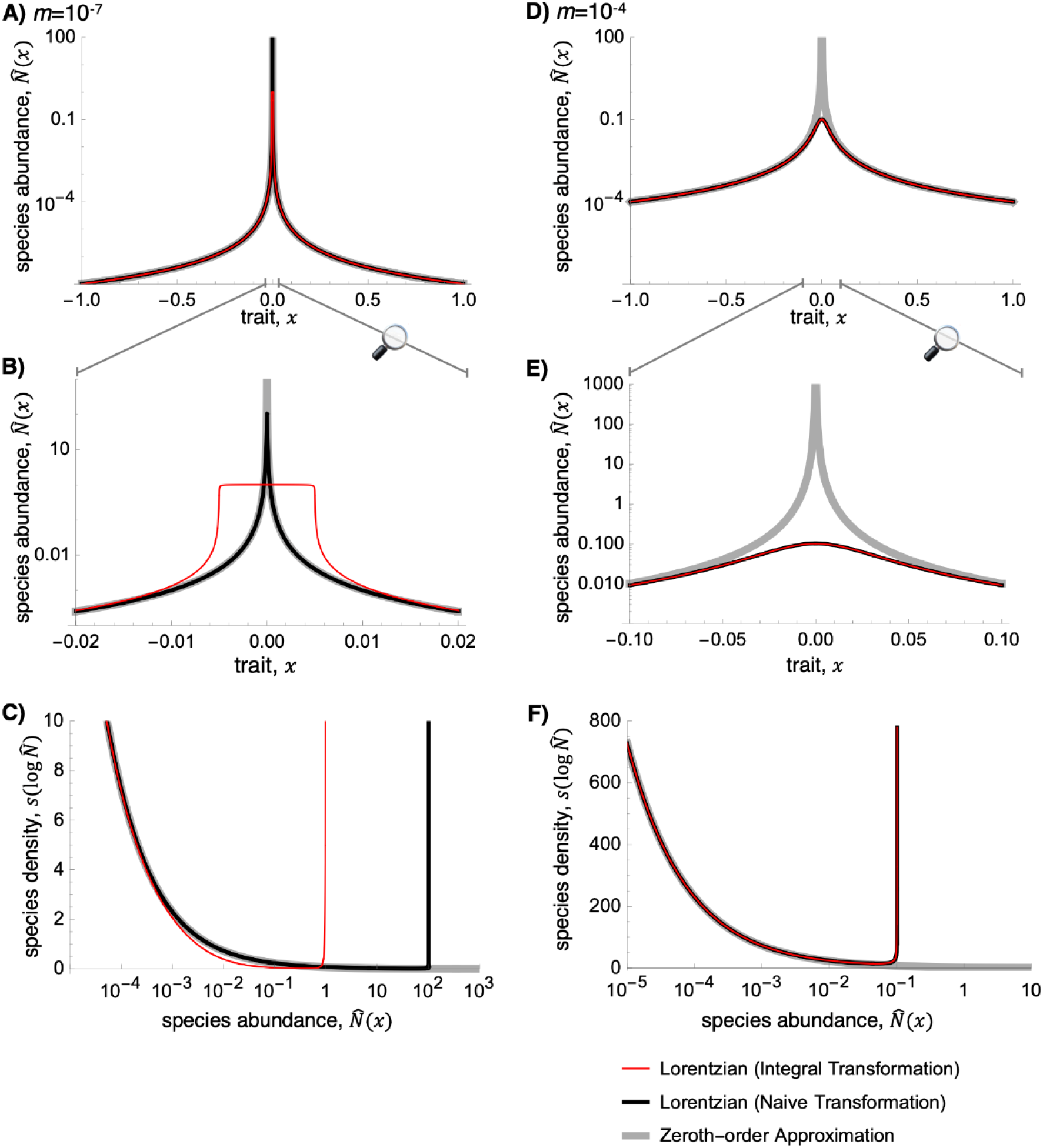
Comparison of **A-B, D-E)** TADs and **C, F)** SADs under various approximations. The thin red line represents the most accurate (but analytically complicated), using the Lorentzian TAD (eqn. 7) along with the integral transformation in Box 1. The black line combines the Lorentzian TAD with the naïve transformation. The thick gray line uses the zeroth-order approximation (eqn. 11), which diverges near the core species. All approaches have the same left-tail behavior. For large dispersal (parts C-F), the naïve transformation works as well as the integral transformation, but for small dispersal (parts A-B) the integral transformation prevents unrealistic abundances of the core species. **A-C)** *m* = 10^−7^, **D-F)** *m* = 10^−4^. Other parameters are as in Fig. 2.

## Appendix D: SAD left-tail behavior details

In this appendix we analytically derive the behavior of the left tail of the SAD (as log*N* → –∞) summarized in eqn. (13) in the main text. As a starting point, we express the density of species s per unit of log-scaled abundance log*N* as:

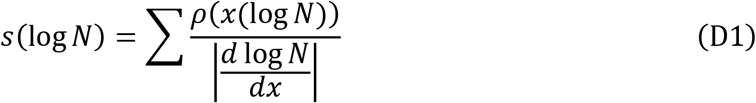

where the sum goes over all the *x*’s associated with log*N*, i.e. all the branches of the (local) inverse function *x*(log*N*) (see Fig. 1). When the inverse function has a single branch, or two symmetrical ones, the sum in eqn. (D1) disappears. In addition, if log*N* (*x*) → –∞ when *x* → ∞, the limit of *s*(log*N*) as log*N* (*x*) → −∞ is

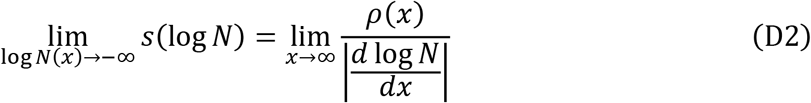

Now, because

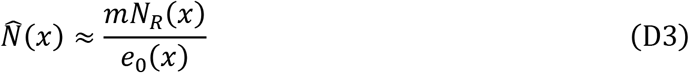

For satellite species with *e*_0_(*x*) ≫ *m* (eqn. (2a)), we have

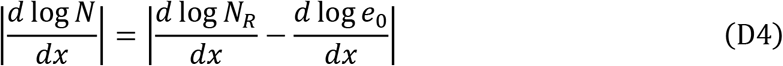

To move further, let us assume as is done in eqn. (12) in the main text that the asymptotic behavior of each function of *x* can be described by a power function times a generalized exponential function; that is,

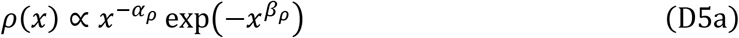

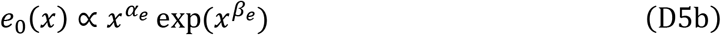

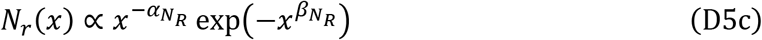

with all exponents non-negative (*α_i_* ≥ 0 and *β_i_* ≥ 0 for *i* = *ρ,e, N_R_*).

Substituting eqn. (D5) into eqn. (D4) we get

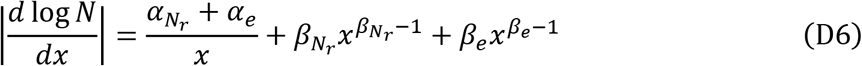

The 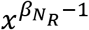 term dominates if *β_N_R__* ≥ *β_e_*, otherwise the *x*^*β_e_*−1^ term dominates if *β_N_R__* < *β_e_*.

Tails of the TADs in these three scenarios can be summarized by

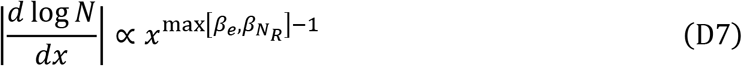

which, when substituted into eqn. (D2), gives the scaling

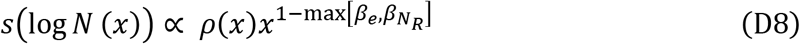

From eqn. (D8) we see that if *ρ*(*x*) decays faster than a power law (*β_ρ_* > 0), then *s*(log*N*) → 0 as *x* → ∞ i.e as *N* → 0. Conversely, if *ρ*(*x*) decays more slowly, as a power law (*β_ρ_* = 0), then:

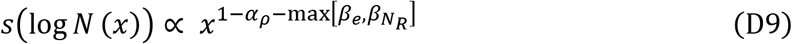

i.e., the left tail of the SAD depends on exponent of the species-density function *ρ*(*x*) relative to the larger exponent of the regional population-density function *N_R_*(*x*) and the exclusion function *e*(*x*):

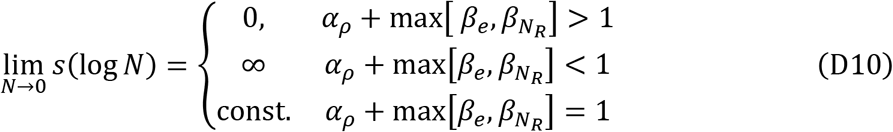

which is eqn. (13) in the main text.

